# Breakdown of Rheological Universality in the Vegetal Hemisphere of Ascidian Embryos Mapped by Atomic Force Microscopy

**DOI:** 10.64898/2026.01.23.701227

**Authors:** Yuki Miyata, Miki Yamamoto, Takahiro Kotani, Yosuke Tsuboyama, Takaharu Okajima

**Author notes:** Correspondence Takaharu Okajima.

## Abstract

Mechanical regulation plays an essential role in the organization of early embryogenesis. In ascidian embryos, cells in the animal hemisphere exhibit periodic stiffening–softening cycles, and their cell rheological properties follow a common master curve, indicating a form of universality. In contrast, cells in the vegetal hemisphere show cell-to-cell differences in stiffness even within the same endodermal lineage, but their rheological behavior has not been characterized. Here, using atomic force microscopy (AFM), we investigated the spatiotemporal dynamics of single-cell power-law rheological states in the vegetal hemisphere during early cleavage. We found that both the elastic modulus (stiffness) and the fluidity (power-law exponent) differ among endodermal cells, and that these differences change in a stage-dependent manner. This result indicates that vegetal cells do not exhibit a single common rheological behavior, in contrast to animal hemisphere cells, suggesting that mechanical properties in the vegetal hemisphere are not uniformly regulated but are patterned in space and time during cleavage. Our findings indicate that this mechanical diversification is linked to the progression of early morphogenesis and may contribute to the emergence of distinct cell behaviors during development.

## 1. Introduction

In embryogenesis, morphology, such as cell positions and cell-to-cell contacts, was highly regulated by mechanical cues ^1–9^. Ascidian embryos have been widely used to investigate how the mechanical state varies during embryogenesis, owing to their simple architecture and invariant early lineages (Fig. 1a and b) ^10–18^. Previous studies revealed that the recruitment and release of actin filaments in the cell cortical region in developing embryos leads to cell stiffening and softening during cell division, exhibiting a periodic mechanical oscillation during embryogenesis ^19,20^. Furthermore, the cell rheology in the animal hemisphere follows a common universal master curve, occupying three discrete mechanical states aligned with cell-cycle phases ^21^. This universality suggests underlying physical constraints on cellular material behavior.

**FIGURE 1.**
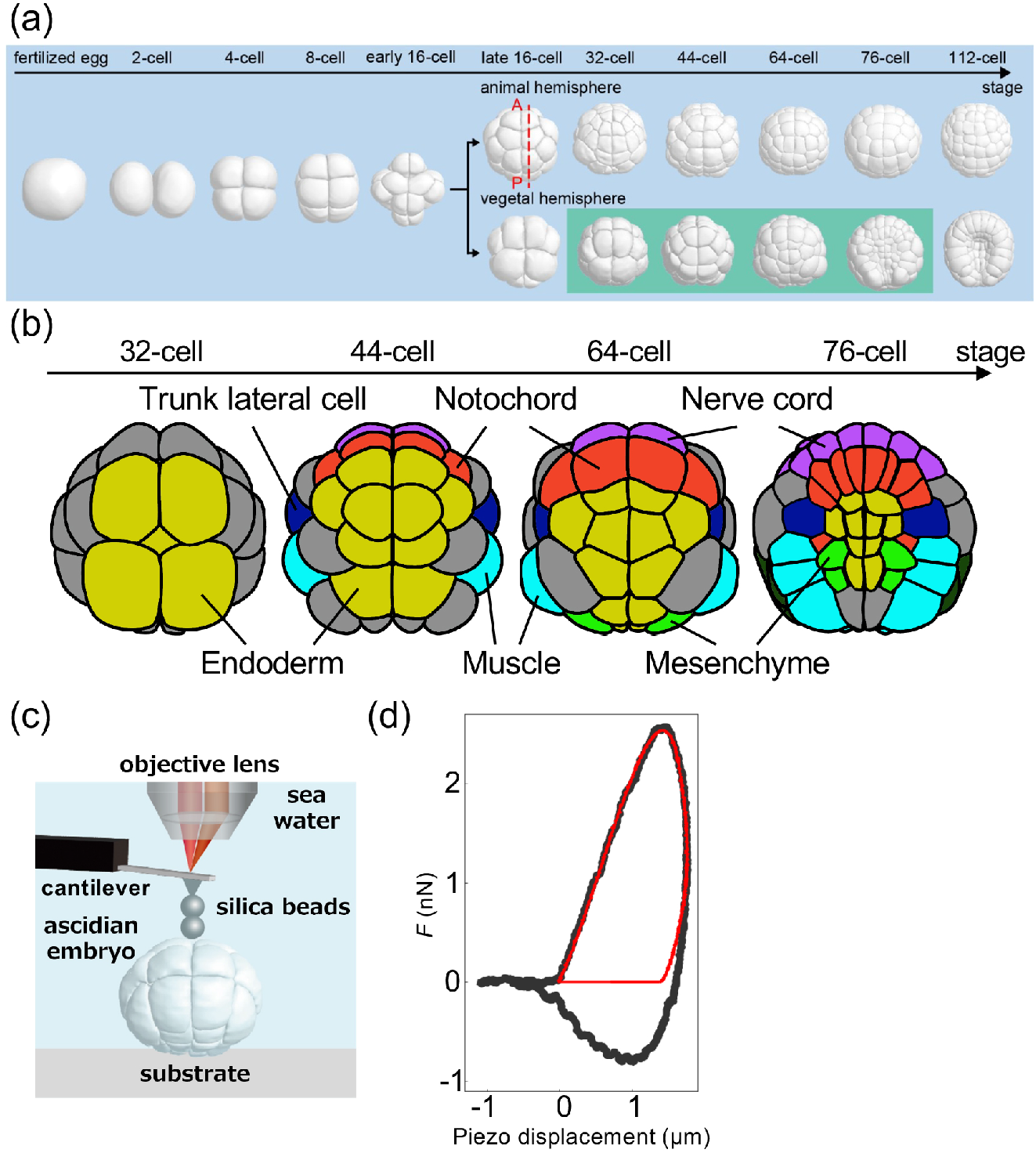
(a) Morphological changes in the 3D ascidian embryo model at early developmental stages from the fertilized egg to the 112-cell stage. Embryos from the 32-cell to the 76-cell stages in the vegetal hemisphere (*light green*) were measured using AFM. The dotted red line represents the anterior-posterior line of the embryo. (b) Morphological changes in cells before cell differentiation, where cell colors represent shared lineages. The colors of dots represent the following cells: ochre (endoderm), red (notochord), light blue (mesenchyme), and purple (nerve chord). (c) Schematic of the AFM setup for measuring developing ascidian embryos weakly adhered to a dish in seawater. (d) A representative AFM approach and retraction force–distance curve acquired from the apical region of the vegetal hemisphere in a developing embryo. The black dots and the red lines represent the measured data and the fitted result with Eqs. 2 and 3.

On the other hand, the stiffness changes show pronounced differences in the vegetal hemisphere ^19^. In the vegetal hemisphere, particularly at the 32- to 44-cell stages, mechanical behavior is more complex. Neighboring pre-endodermal cells, such as A6.1 and B6.1 cells, stiffen synchronously before mitosis, resembling the animal hemisphere. Yet, their post-division trajectories are separated into two mechanical states: A7.1/A7.2 cells, which are divided from the A6.1 cell, soften rapidly after cytokinesis, whereas B7.1/B7.2 cells, divided from the B6.1 cell, maintain elevated stiffness into later stages approaching gastrulation. These observations suggest lineage-dependent mechanical regulation, with vegetal cells maintaining stiffness that is not simply tied to the mitotic cycle. However, the rheological properties of cells in the vegetal hemisphere remain unknown.

Here, using atomic force microscopy (AFM) ^22,23^, we map the spatiotemporal evolution of cell rheology in the vegetal hemisphere across early cleavage. Our observations indicated a breakdown of rheological universality observed in the animal hemisphere, indicating that early embryonic mechanics are not uniformly regulated but rather follow spatiotemporally organized rules, and that the loss of rheological universality provides a physical framework linking cell-level material states to the coordinated progression of morphogenesis.

## 2. Materials and Methods

### 2.1 Ascidian samples

Adults of *Ciona intestinalis* type A (*Ciona robusta*) were collected near the Maizuru Fisheries Research Station (Kyoto University) and the Misaki Marine Biological Station (The University of Tokyo) via the National BioResource Project (NBRP, Japan). Gametes were obtained by dissecting the gonoducts. Unfertilized eggs were dechorionated in natural seawater (Nihon Aquarium) containing 1.0% sodium thioglycolate (Wako Pure Chemical Industries, Ltd.) and 0.05% Actinase E (Rikaken Pharmaceutical Co., Ltd.). After several washes in seawater, eggs were fertilized in vitro and cultured on 0.1% gelatin–coated dishes (Wako Pure Chemical Industries, Ltd.) in seawater at 18 °C. For AFM measurements, embryos were gently immobilized on culture dishes (Iwaki) in seawater to minimize lateral drift during recording. Embryos that failed to develop normally at early stages were excluded; only embryos that progressed to the gastrula stage were analyzed.

### 2.2 AFM measurements

A customized atomic force microscope ^21,24^ was used to map the relative height and rheological properties of ascidian embryo samples. The atomic force microscope was mounted on an upright optical microscope (Eclipse FN1; Nikon) with a liquid-immersion objective lens (plan fluor, ×10, Nikon) for the optical lever system, as reported previously 19-21,24. We used a rectangular cantilever (Biolever-mini, BL-AC40TS-C2; Olympus) with a nominal spring constant of < 0.1 N/m. Before the experiments, the spring constant of the cantilever was determined in liquid environments using a thermal fluctuation method. To achieve a well-defined contact geometry and prevent the contact between the cantilever beam and the sample surface, two silica beads with a radius *R* of ca. 5 μm (Funakoshi) were arranged tandemly from the AFM tip with epoxy glue (Nichiban) (Fig. 1c). The AFM force mapping measurements were conducted in a part of the vegetal hemisphere of the embryo at 18°C with a piezo-scanner (E-761; Physik Instruments) controlled with LabVIEW software (National Instruments). The embryo samples were weakly immobilized on culture dishes, so that abnormal cell division was highly prevented. However, the weak adhesion occasionally caused small, unexpected fluctuations. Thus, the AFM-mapped positions were manually adjusted to trace approximately the same region of the embryo throughout development, accounting for sample fluctuations. The scan range was about 72 μm × 72 μm with a spacing of 3 μm. The acquisition time for a single mapping image was approximately 3 min at a slow scan speed, reducing the risk of unexpected movement of embryo samples. The cells mapped by AFM were identified using the Four-Dimensional Ascidian Body Atlas database ^14^.

### 2.3 AFM analysis

The time-dependent relaxation modulus *E*(*t*) of a power-law rheology (PLR) model ^23,25-^ 30 is expressed as:

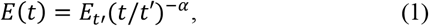

where *E*_*t’*_ is the modulus scaling parameter at time *t’*, which corresponds to the relaxation modulus *E*(_*t’*_), and *α* is the power law exponent. According to the soft glassy rheology model ^26,31^, *α* is an effective temperature that represents the magnitude of agitation within an energy landscape of material configurations containing many wells or traps, and rheological materials exhibit an *α* value between a complete solid (*α* = 0) and a fluid (*α* = 1). Thus, *α* is referred to as the “fluidity” ^25,26,31^.

Using the tilt angle *θ* of the sample surface, the approach and retraction force-indentation curves were analyzed using Ting’s model ^23,28,32^ with the modified Hertz contact model ^33^. The loading force *F* is given by

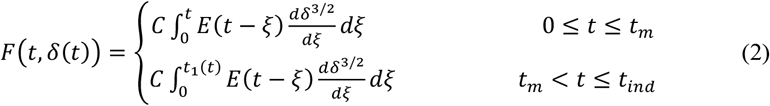

and

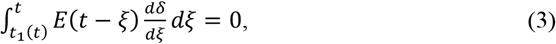

where *C* = (4*R*^1/2^ cos^5/2^*θ*/[3(1-*ν*_p_^2^]) is the geometrical pre-factor for a spherical probe with a radius of *R*, and *δ* is the indentation depth. The Poisson’s ratio of the cell, *ν*_p_, was assumed to be 0.5. *t*_m_ is the duration of the approach curve, *t*_ind_ is the duration of the approach and retraction curves, and *t*1 is the auxiliary function determined using Eq. 3 (Fig. 1d). The *θ* was estimated from the corresponding *H* image. We discussed the phenomena observed in mapping regions that satisfied θ < 40°, because the modified Hertz contact model was valid for *θ* < 40°^33^. The modulus scaling factor *E*_t_’ at time *t*’ maps were log-scaled using *ν* = log_10_*E*_t_’ (Pa), whereas the *H* maps were presented on a linear scale. For data analysis, an unpaired two-tailed Student’s *t*-test was used to determine *p*-values. The *p*-values less than 0.05 were significant.

## 3. Results

### 3.1 Rheological changes of the ascidian embryo from the 32-cell to the 44-cell stage

We measured spatiotemporal rheological properties of the ascidian embryo in the vegetal hemisphere during early cleavage from the 32-cell to 76-cell stage (Fig. 2a). At the 32-cell stage, we measured the rheological properties of A6.1 and B6.1 cells, which are pre-endodermal. As a result, the modulus scaling factor at *t’* = 1 s, *E*_1_, was higher in A6.1 cells compared to B6.1 cells, whereas the power-law exponent, α, was significantly lower in A6.1 than in B6.1 cells (Fig. 2b).

**FIGURE 2.**
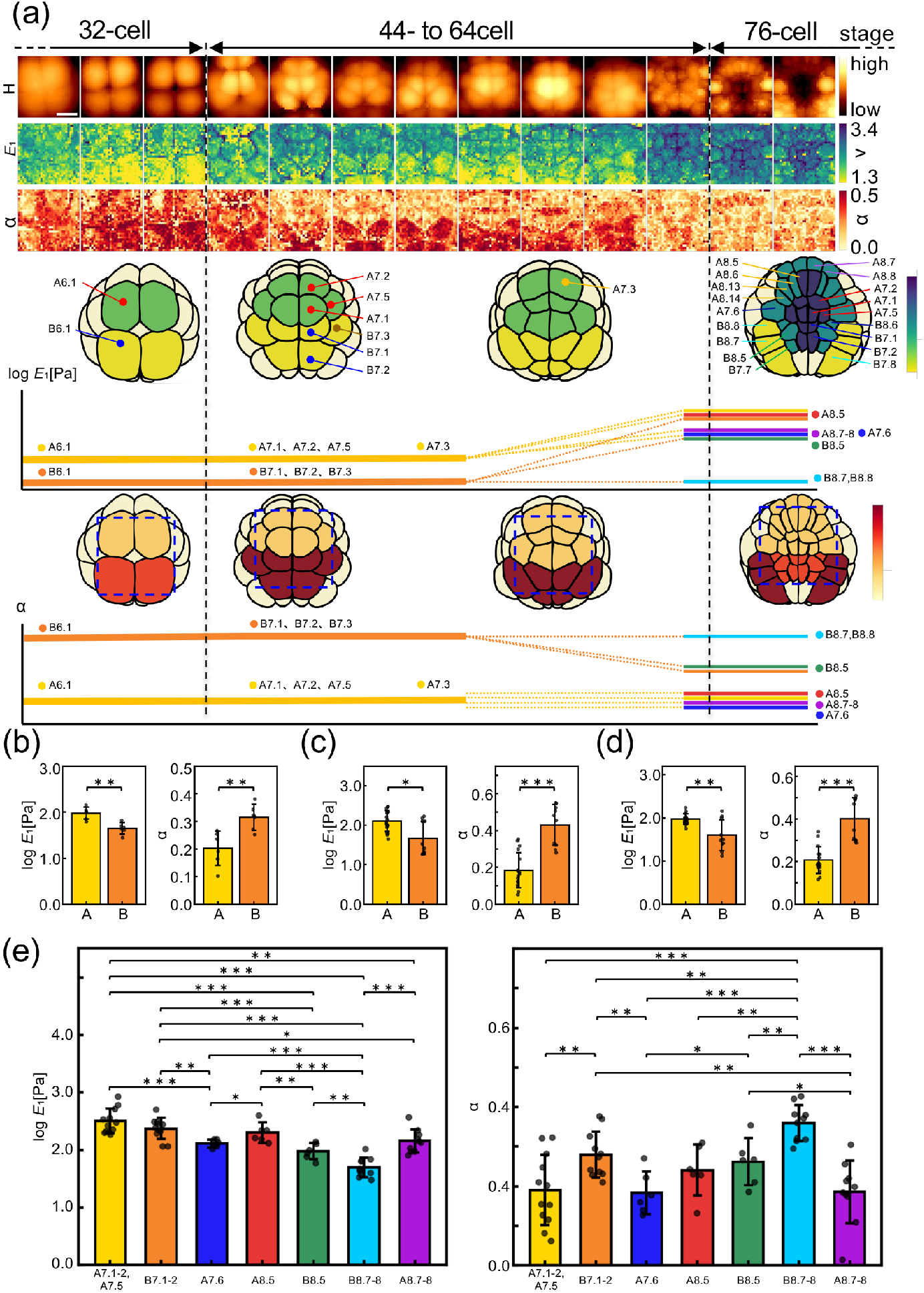
(a) Portions of *H, E*_1_, and *α* images mapped in the embryo in the vegetal hemisphere from the 32-cell to the 76-cell stages using the approach-retraction AFM force curve measurement. Scale bar was 30 μm. Temporal changes in rheological parameters quantified from (b) – (e) were schematically summarized in a 3D model with colors. (b) Modulus scaling parameter at 1 s, *E*_1_(geometric mean± s.d.) and power-law exponent, α (arithmetic mean± s.d.), of A6.1 (denoted as A) and B6.1 (denoted as B) in the 32-cell stage from *n* = 3 embryos. *E*1 (geometric mean± s.d.) and α (arithmetic mean± s.d.), of A7.1/A7.2 (denoted as A) and B7.1/B7.2 (denoted as B) in the 44-cell stage (c), the 64-cell stage (d) from *n* = 3 embryos. (e) *E*_1_ (geometric mean± s.d.) and α (arithmetic mean±s.d.) of different pre-determined cells before cell differentiation in the 76-cell stage. \**p* < 0.05. \*\**p* < 0.01.\*\*\**p* < 0.001

During the 32-cell to 44-cell stages, embryonic cells A6.1 and B6.1 divide into A7.1/A7.2 and B7.1/B7.2, respectively (Fig. 2a). We observed that *E*_1_ was consistently higher in A7.1/A7.2 cells than in B7.1/B7.2 cells. In contrast, α in A7.1/A7.2 cells was lower than that in B7.1/B7.2 cells (Fig. 2c). Proceeding the developmental stages from the 44-cell to the 64-cell stage, the same significant difference in both *E*_1_ and *α* was observed (Fig. 2a and d).

### 3.2 Rheological changes of the ascidian embryo from the 66-cell to the 76-cell stage

From the 64-cell stage—when cell fates are effectively predetermined ^34,35^—the geometry of the ascidian embryo enables lineage-resolved AFM mapping, allowing us to probe the mechanical properties of distinct fate-specified cells in vivo. Upon entry into the 76-cell stage, the rheological properties of cells measured by AFM were distributed across three different states. In the *E*_1_ value, pre-endoderm cells (A7.1/A7.2/A7/5 and B7.1/B7.2) with A8.5 (pre-notochord) cells became the highest level. The intermediate level was A7.6 (trunk lateral) cells, A8.7-8 (pre-neuronal) cells, and B8.5 (pre-mesenchyme) cells. The lowest level was B8.7-8 (pre-tail muscle) cells (Fig. 2a and e).

In parallel, the *α* values measured by AFM also occupy three states. The highest level was B8.7-8 (pre-tail muscle) cells. The intermediate level was B7.1/B7.2 (pre-endoderm), B8.5 (pre-mesenchyme), A8.5 (pre-notochord), and B8.6 (pre-notochord) cells. The lowest level was A7.1/A7.2/A7/5 (pre-endoderm), A8.5 (pre-notochord), A8.7-8 (pre-neuronal), and A7.6 (trunk lateral) cells (Fig. 2a and e).These results indicated that the changes in power-law rheological parameters differed among cells with distinct fates before cell differentiation. The time evolution of cell rheology observed in the vegetal hemisphere is summarized schematically in Fig. 2a.

## 4. Discussion

### 4.1 Power-law rheology of ascidian embryo measured by AFM

The AFM maps of *E*_1_ and *α* reproduced cell morphologies and boundaries observed in the relative height, *H*, image, and the positions of individual cells matched those assigned by the confocal microscopy–based 3D virtual embryo model (Fig. 1b). This agreement confirms that the rheological parameters of cells in the vegetal region extracted from AFM force curves can be attributed to the cells directly contacted by the AFM probe. In ascidian embryos, previous AFM work further suggested that such measurements predominantly probe cortical actomyosin structures rather than the deep cytoplasmic interior, supporting the interpretation that our maps reflect the mechanical state of the apical cortex.

Consistent with a cortical origin, pharmacological perturbations reported previously showed that interphase stiffening in the embryo—occurring without cell division—was markedly reduced by inhibition of the Rho kinase (ROCK) pathway using Y-27632 ^19^. In addition, immunofluorescence analyses have shown that ROCK inhibition relaxes apical shrinkage in the vegetal hemisphere and reduces apical and circum-apical myosin accumulation ^36^. Together, these observations suggest that the Rho–ROCK pathway modulates the rheological properties measured by AFM, through regulating myosin enrichment at the apical region of vegetal cells ^1,12,36^.

### 4.2 Breakdown of Rheological Universality

Cell rheology is increasingly recognized as a key physical factor in morphogenesis and developmental progression ^37–40^. In isolated single cells in vitro, diverse perturbations of intracellular structures often cause the PLR parameters to collapse onto a common master curve, ^25–27,41–47^ although the physical origin of this relationship remains unresolved. In the context of the ascidian embryo, AFM mapping previously revealed that embryonic cells can occupy discrete power-law rheological states across the cell cycle, where stiffness and fluidity exhibit a negative correlation and collapse onto a master curve. These findings imply that cells embedded in the embryo can maintain universal rheological behavior comparable to that observed in isolated cells.

To examine whether a similar correlation holds for vegetal-hemisphere cells during early cleavage, we plotted embryo-level means of *E*_1_ (geometric mean) and α (arithmetic mean) for the embryos shown in Fig. 3a. At the 32-cell stage, both A6.1 and B6.1 cells aligned on rheological master curve #1 (MC #1) (Fig. 3a and b), which is identical to that reported for the animal hemisphere. After division at the 44-cell stage, the daughter cells A7.1 and A7.2 showed decreased *α* and increased *E*_1_ while remaining on MC #1. In contrast, B7.1 and B7.2 increased both *E*_1_ and *α*, deviating from MC #1 and shifting toward a distinct master curve #2, MC #2 (Fig. 3a and b). At the 64-cell stage, A7.1/A7.2 decreased *E*_1_ and increased α while still following MC #1, whereas B7.1/7.2 showed little change and maintained MC #2 (Fig. 3a and b). Upon reaching the 76-cell stage, A7.1/A7.2 increased both *E*_1_ and *α* and transitioned to MC #2, while B7.1/B7.2 gradually increased *E*_1_ and decreased *α* along MC #2 (Fig. 3a and b). Consequently, A7.1/A7.2 converged onto the same master curve as B7.1/7.2. These observations indicate that, even among pre-endodermal lineages sharing a common fate, rheological trajectories can diverge before gastrulation and later converge, implying a temporally structured regulation of mechanical states in the vegetal hemisphere.

**FIGURE 3.**
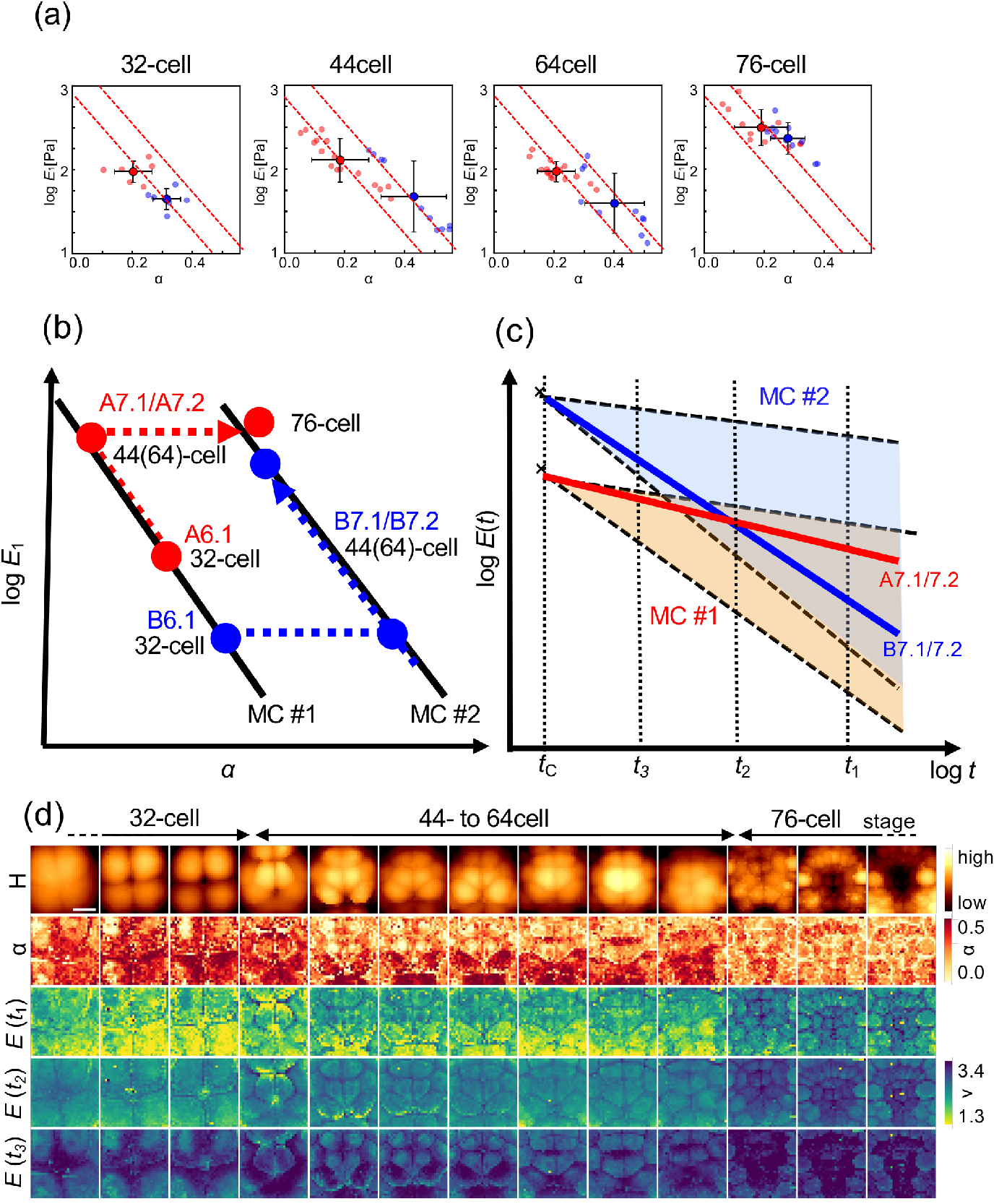
(a) Plots of *E*_1_ (geometric mean± s.d.) versus *α* (arithmetic mean± s.d.) of pre-endodermal (A6.1 in red dots and B6.1 in blue dots in the 32-cell stage, A7.1/A7.2 in red dots and B7.1/B7.2 in blue dots during the 44-cell to 76-cell stages) in rheological states for embryos (*n* = 3) where the rheological parameter of each endodermal cells were also plotted. The slope of the left-sided red dotted line was similar to that previously observed in the animal hemisphere of the ascidian embryo at early stage ^21^. (b) Schematic picture of temporal changes of rheological parameters of endodermal cells on two master curves observed in (a). (c) Schematic picture of relaxation modulus of endodermal cells as a function of time on different master curves during the 44-cell to 64-cell stages from (b). The red and blue relaxation curves, corresponding to A cells and B cells, respectively, can change within their respective regions fixed at the crosspoint at a short timescale of *t*_3_. (d) The AFM modulus scaling factor images obtained in Fig. 2a were mapped using different timescales: *t*_1_=1 s, *t*_2_=10^−2^ s, and *t*_3_=10^−4^ s, together with *H* and *α* images shown in Fig. 2a.

### 4.3 Relationship between the apparent Young’s modulus and Rheological Parameters

Previous AFM measurements further reported that during the 44- to 64-cell stages, the observed Young’s modulus of B7.1/B7.2 cells exceeds that of A7.1/A7.2 cells, suggesting that B-line cells are stiffer. In contrast, our PLR analysis indicates that the modulus scaling parameter *E*_1_ can be higher in A7.1/A7.2 than in B7.1/B7.2 at the same stages. This apparent discrepancy can be reconciled if A7.1/A7.2 and B7.1/B7.2 occupy distinct rheological master curves with different *α*. In case the rheological parameters follow a master curve, the relaxation modulus *E*(*t*) crosses at a higher frequency when changing the rheological parameters ^25–27,41,42,47^. Based on Fig. 3b, we can schematically describe the relaxation modulus of A7.1/7.2 and B7.1/7.2 cells in the 44 to the 64 cell stage (Fig. 3c).

Two relaxation responses corresponding to MC #1 and MC #2 cross depending on the evaluation timescale (Fig. 3c). The modulus scaling parameter *E*(*t*_3_) at a short timescale *t*_3_ increases so that B7.1/B7.2 can appear stiffer than A7.1/A7.2. The B7.1/B7.2 is comparable to A7.1/A7.2 in the intermediate time scale *t*_2_, where the relaxation modulus crosses, and B7.1/B7.2 can appear softer than A7.1/A7.2 in a long timescale *t*_1_. This interpretation is supported by modulus scaling parameter maps generated with different timescale factors, which show a timescale (*t*_2_ = 10^−2^ s) at which the apparent modulus of A7.1/A7.2 and B7.1/B7.2 converges (Fig. 3d). Therefore, comparisons of stiffness across lineages must be interpreted in the context of the characteristic timescale of the measurement. Master-curve transitions provide a unifying framework for seemingly conflicting stiffness readouts.

## 5. Conclusions

In this study, we performed lineage-resolved AFM mapping of power-law rheology in the vegetal hemisphere of ascidian embryos during early cleavage. By quantifying both the modulus scaling parameter at 1 s, *E*_1_, and the power-law exponent (fluidity), *α*, we revealed stage-dependent and cell-to-cell differences in rheological states within pre-endodermal lineages. These results demonstrate that, unlike cells in the animal hemisphere that collapse onto a single universal master curve, vegetal cells follow distinct rheological master-curve trajectories and can transition between rheological states as development proceeds. Furthermore, our analysis provides a timescale-based framework to reconcile apparent discrepancies in stiffness readouts obtained at different measurement timescales, emphasizing that lineage comparisons depend on the measurement timescale. Together, our findings show that mechanical properties in the vegetal hemisphere are spatiotemporally patterned during cleavage and suggest that early mechanical diversification may contribute to the emergence of distinct lineage behaviors during progression toward gastrulation.

## Acknowledgments

We thank the National BioResource Project (NBRP, Japan) for providing *C. intestinalis* samples from Dr. Yutaka Satou (Kyoto University) and Dr. Manabu Yoshida (The University of Tokyo). We also thank Dr. Kohji Hotta (Keio University) for advising the sample preparation method and Dr. Kaori Kuribayashi-Shigetomi (Hokkaido University) for assistance with editing the manuscript.

## Author contributions

T.O. conceived and designed the experiments. Y.M., M.Y., T.K., and Y.T. performed AFM experiments and the data analysis. Y.M., M.Y., and T.O. prepared the manuscript. All authors edited the manuscript before submission.

